# A framework for quantifying the mechanics of dexterous grasp

**DOI:** 10.64898/2026.05.05.723084

**Authors:** Anton R. Sobinov, Xuan Ma, Elizaveta V. Okorokova, Charles M. Greenspon, Caleb Raman, Natalya Shelchkova, Qinpu He, Neema Darabi, Rashi Bhatt, Patrick Jordan, Paul Arters, Nicholas G. Hatsopoulos, Lee E Miller, Sliman J. Bensmaia

**Affiliations:** Department of Organismal Biology and Anatomy, University of Chicago, Chicago, IL, USA; Department of Neuroscience, Northwestern University, Chicago, IL, USA; Department of Neurological Surgery, University of California, Davis, CA, USA; Department of Neurological Surgery, University of Chicago, Chicago, IL, USA; Department of Neurology, University of Chicago, Chicago, IL, USA; Neuroscience Institute, University of Chicago, Chicago, IL, USA; Department of Biomedical Engineering, Northwestern University, Evanston, IL, USA; Department of Physical Medicine and Rehabilitation, Northwestern University, Chicago, IL, USA

## Abstract

A hallmark of primate behavior is the exceptional ability to dexterously grasp and manipulate objects, yet the investigation of the neural mechanisms that support manual dexterity has been hindered by technical challenges. Optical hand tracking is complicated by frequent occlusions, and contact forces are hard to measure with sufficient precision. Furthermore, while monitoring the kinematics during reaching phase, and the contact forces during object manipulation phase is difficult, joint torques are impossible to measure directly. While challenging, the ability to estimate joint torques in the complex primate hand could provide an invaluable continuous mechanical description spanning both phases. With this in mind, we have developed an experimental apparatus and data processing pipeline for quantifying these variables describing prehension. The apparatus presented objects of various sizes and orientations throughout the workspace, evoking different grasping strategies. Object surfaces were instrumented with thousands of pressure-sensitive elements, enabling high-resolution measurement of distributed contact forces. Simultaneously, eight high-speed cameras were used to reconstruct hand and arm movements with markerless tracking, triangulating 3D landmarks, and mapping them onto a musculoskeletal model, enabling estimation of time-varying joint angles. This posture quantification allowed contact forces to be automatically assigned to specific hand segments, in close agreement with manual human annotations. We used the reconstructed movements and contact forces with the musculoskeletal model of the hand to compute inverse dynamics, yielding joint torques throughout behavior, unifying the hand kinematics and grasp forces into a single physical description. Throughout the processing, we identified individual neurons in the motor cortex of monkeys that were related to grasp force, kinematics, and torques. Together, this framework enabled a comprehensive and precise physical characterization of primate manual behavior, providing a foundation for investigating the neural mechanisms of manual dexterity.

## Introduction

Grasping and manipulating objects is a hallmark of primate behavior, particularly humans. We can position and shape our hand to grasp an object optimally based on its shape, and then apply a complex symphony of forces with different parts of the hand to manipulate it (Castiello, 2005; Johansson and Flanagan, 2009). An expansive neural apparatus unique to primates supports this behavior (Dum and Strick, 2002; Hinkley et al., 2010; O’Connor et al., 2021; Sobinov and Bensmaia, 2021). Recent work on hand control in reduced behavioral spaces has been especially useful for testing and refining theories of motor control (Churchland et al., 2012; Gallego et al., 2017; Natraj et al., 2022). Natural object manipulation, by contrast, involves much greater variation in posture and interaction forces (Yan et al., 2020). However, measuring or computing these rapidly changing high-dimensional signals remains a formidable technical challenge (Jiang et al., 2024; Jin et al., 2024).

Prehension – the act of reaching toward and grasping an object – comprises two aspects: moving the hand in space while shaping it, then applying forces with the digits and palm to the object (**Figure 1**) (Jeannerod, 1984; Castiello, 2005). Although the kinematics of prehension and associated neural activity have been studied extensively, investigation of the associated interaction forces has been limited by the difficulty collecting these data (Jakobson and Goodale, 1991; Fluet et al., 2010; Townsend et al., 2011; Yan et al., 2022). Between the skin and an object arise many points of contact force, such as those exerted by individual finger pads during grasp or by a single digit when pressing a keyboard key. In principle, these forces can be measured using sensors mounted on the object, but tracking the specific skin region making contact becomes difficult when the object or sensor shifts relative to the skin or when the grasp changes. These forces evoke activity throughout the somatosensory neuraxis and providing feedback that is critical to manual behavior (Hermsdörfer et al., 2008; Johansson and Flanagan, 2009; Yau et al., 2016; Delhaye et al., 2018; Parry et al., 2021; Suresh et al., 2021).

**Figure 1.**
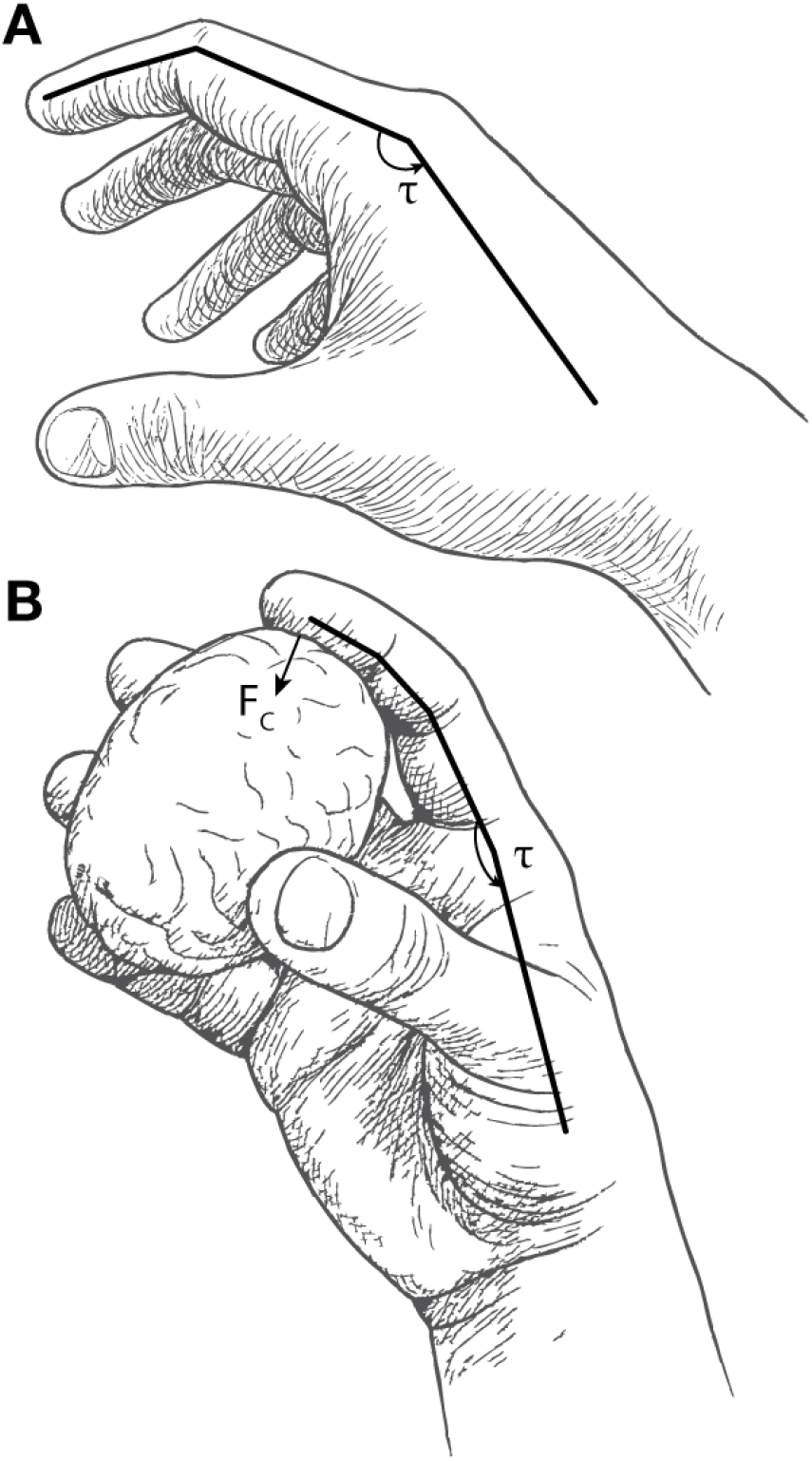
Aspects of prehension with a schematic representation of forces. **A**| Free movement of digits. **B**| Manipulating an object. τ – metacarpophalangeal actuating torque; F_c_ – contact force between index finger digit pad and the object.

Contact forces during manipulation arise from the actuating forces produced by muscle contraction, combined with the effects of tendons and other passive elements. In biomechanical analyses, these effects are often summarized as net torques acting about individual degrees of freedom of a joint, although these studies have mostly focused on the proximal joints – shoulder and arm, where modeling is somewhat more tractable, particularly if action is restricted to a horizontal plane, allowing gravity to be ignored (Flash and Hogan, 1985). These simplifications do not readily extend to the hand, whose many small segments, coupled tendons, and multiarticular muscles make estimation of joint torques substantially more difficult (Valero-Cuevas, 2005; Binder-Markey and Murray, 2017; Valero-Cuevas and Santello, 2017).

By definition, contact forces are absent outside periods of object contact, and vary dramatically upon contact (Santello and Soechting, 1998; Johansson and Flanagan, 2009; Yan et al., 2026). In contrast, hand kinematics vary throughout reach and preshaping, and remain relatively stable once grasp is established. Joint torques, though difficult to compute, might bridge these two phases of prehension, providing a continuous mechanical description of the behavior. In this manuscript, we describe a method for tracking contact forces and kinematics, while modelling joint torques, as monkeys grasped various objects. We then combine these measurements with single neuron recordings to illustrate analyses that could generalize to a broad range of electrophysiological datasets.

## Methods and Results

The approach and results we report here outline the experimental apparatus we designed to collect raw behavioral data, the transformation of that raw data into joint angles, and the inference of the forces and torques exerted by each segment of the hand, during its interaction with the object. Ultimately, we are also able to calculate the actuating torques, which are inaccessible to direct measurement. Throughout the manuscript, we provide examples of the neural data collected in the experiment and its relation to the behavioral data.

The monkeys were trained to reach, grasp, and hold an object. In a typical trial, the width and orientation of the object were visible to the monkey, and the force it needed to match was displayed on a multi-segment LED strip (**Figure 2**A). The object was brought to the monkey and a “go” cue sounded, prompting the monkey to reach for and grasp the object. If the target force was achieved and held for a set period of time, a reward was provided. Otherwise, a “fail” tone was played. In either case, the object was withdrawn to prepare for the next trial.

**Figure 2.**
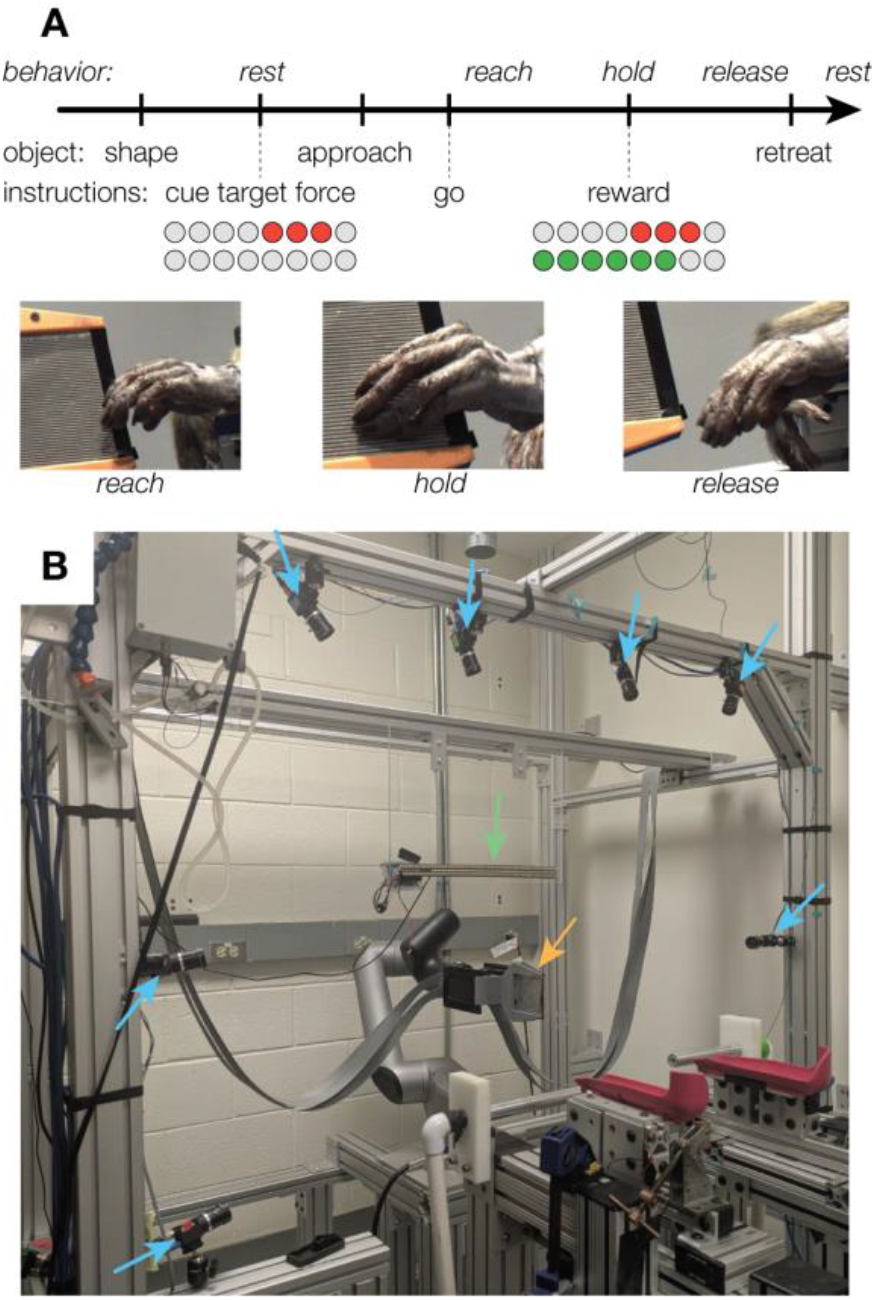
Experimental progression and apparatus. **A**| Typical trial progression. Behavior describes what the monkey is doing, object – how the robot is moving the object, instructions – instructions to the monkey. **B**| The rig actuated by a robotic arm. The location of the cameras is indicated by blue arrows, object instrumented with pressure sensors – orange, LEDs – green. One camera is obscured by the arm rest.

### Experimental apparatus

The experimental apparatus consisted of an actuating system, a recording system, and a feedback system (**Figure 2**B). The actuating system positioned a configurable object in front of the monkey. The recording system, split between several computers, recorded videos from 8 cameras (blue arrows around the periphery), high-resolution force from the pressure sensors mounted on the object (orange arrow), and electrophysiological data. The real-time feedback system provided the monkey with information about force exerted on the object (green arrow). A single program, controlled by the experimenter, communicated with the separate recording programs, set task and device configurations, and synchronized recording. It first sent a synchronization pulse to all recording systems, and then triggered the start of pressure sensor and camera recording (**Supplementary Figure 3**).

The system can generate a variety of objects from two rigid parallel aluminum plates whose aperture and orientation were varied across trials (orientation range −90° to 90°, typically −30°, 0°, or 30°; object width 1.0–2.5 cm). To reduce electrical noise coupling from actuation into plate-mounted sensors, the plates were electrically insulated from the frame and sensor cabling was mechanically secured and shielded.

We developed two robotic systems to position the grasp object throughout the monkey’s peripersonal workspace (**Figure 2**B) with a programmable apparatus. One used servo-driven stages to translate the object, adjust tilt and aperture, with motors continuously engaged to prevent unintended motion (**Supplementary Figure 4**). For more flexible positioning, we used an industrial robotic arm (Kinova Link 6) and gripper (OnRobot 2FG7) carrying the same plate attachments, controlled via Kinova’s Python API using endpoint position commands for both the arm and gripper (displayed in **Figure 2**B). The increase in degrees of freedom came with an increase in complexity of control, since all target locations of the object had to be precomputed by the robot using a built-in inverse kinematic system.

We used two interchangeable sensor approaches to enforce consistent initial and final rest posture of the monkey. In the first one, light-sensitive photoresistors in the arm rest aborted trials if the hand moved early. Trial completion was registered only after the arm returned to the rest posture, with an extra reward to encourage full sequences. In the alternative approach, conductive metal handlebars that detected grasp contact helped avoid sensitivity to ambient light and hair occlusion.

#### Force measurements

We sought to measure not only total grasp force but also its distribution across the digits and individual finger pads (**Figure 3**A). Each plate comprising the object was instrumented with a high-density pressure sensor (Tekscan 5076; 44×44 sensels over 84×84 mm), providing 4-16 sensels per digit segment (e.g., ∼13 sensels across the distal phalanx of the index finger of a monkey). Early in the experiments, the monkey occasionally grasped the object from the top or bottom, where portions of the array were unsensorized. To encourage consistent contact with the central sensorized region, trial success was based on total force computed from a masked subset of sensels, excluding the top and bottom 10% of the array. Data were recorded continuously at 300 Hz, with a parallel 60 Hz stream used for real-time display and force feedback, the rate chosen to avoid frame drops. Sensors were equilibrated and calibrated before mounting using an air-bladder procedure to apply equal pressure to all sensels. For subsequent calibration, we applied known loads wrapped in latex gloves to mimic the experimental contact interface.

**Figure 3.**
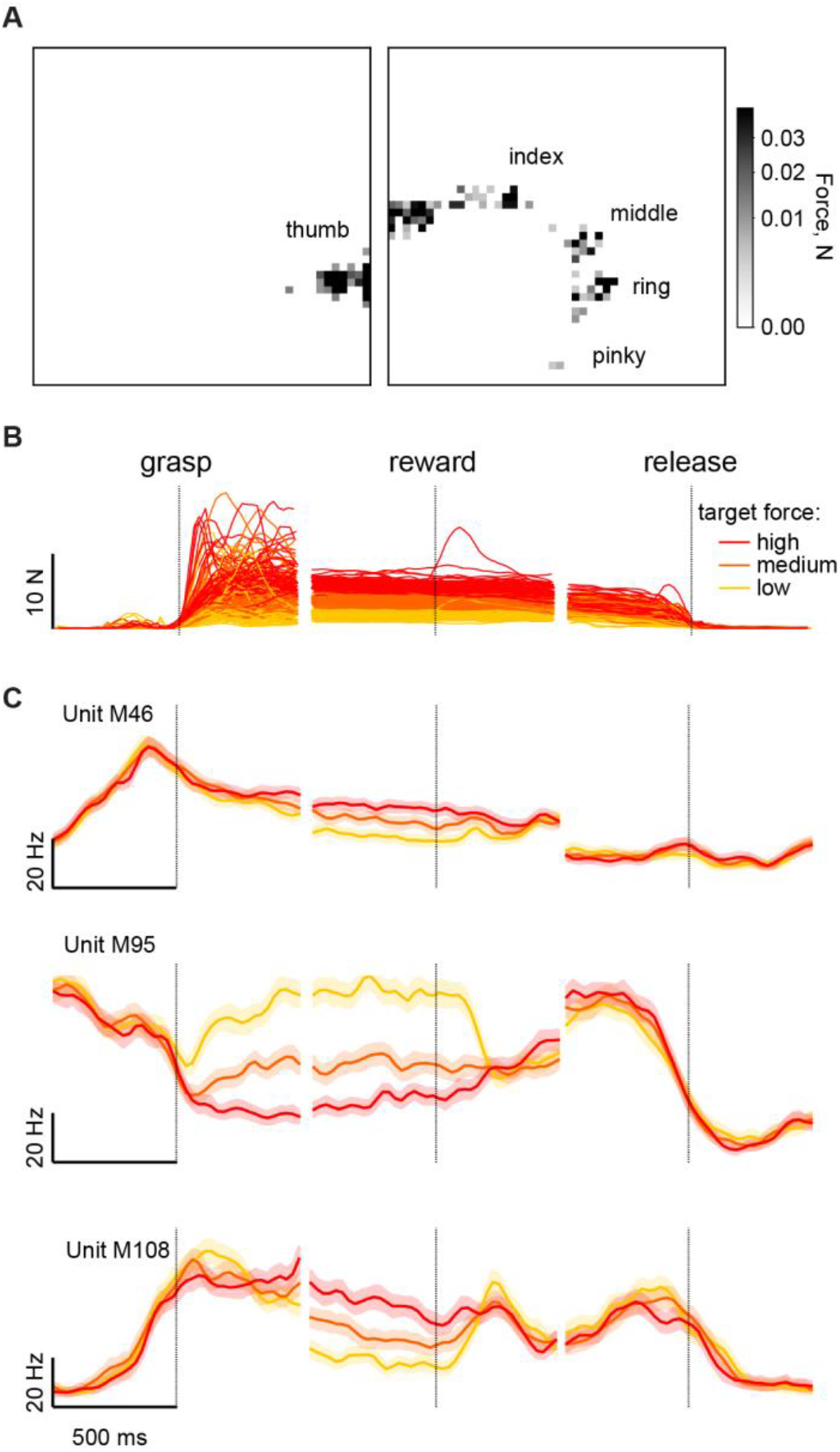
Pressure sensor data. **A**| A snapshot of the pressure maps on both sides of the grasped object. A dark spot on the left sensor corresponds to the thumb tip, a line along the edge – the thenar part of the hand touching the sensors. The four spots on the right are the tips of four fingers, and the spot near the proximal edge – the base of the index finger. **B**| Total force exerted on both plates during a single session with monkey M aligned to three time points – initial grasp, first reward administration, and release. The color of the line corresponds to three levels of target force. **C**| The PETHs of three well isolated M1 units from the same session of monkey M. The activity of the units is aligned to the same three time points. The mean (solid lines) and standard errors (shaded areas) are shown for ∼80 trials per force level.

After each session, pressure data were converted from the proprietary Tekscan format into a time-indexed tensor of forces for simpler and faster access (Sobinov, 2024). We converted the pressure measured by each sensel into force by multiplying it by the area covered by a sensel. Force signals were processed to remove electrical noise and static offsets caused by sensor deformation (e.g., protective sheath pinching some of the sensors). We defined “active” periods as those in which the total force exceeded 5% of the maximum total force. We computed a threshold map indicating the 90^th^ percentile of the force recorded by each sensel outside of the active period. Values below threshold were zeroed, and the signals filtered with a 10 ms median filter. There was no spatial filtering. This produced stable force traces without electrical noise and deformations across the entire session.

Throughout a single session, monkeys were tasked to exert forces within specified bounds around several target forces (**Figure 3**B). We observed different modulation profiles of motor cortical neurons associated with the grasp force (**Figure 3**C). Some were positively correlated with force (e.g., M46), and some had inverse correlations (e.g., M95). Some even changed the relationship throughout the trial (e.g., M108 from no modulation to positive correlation).

#### Cameras

We recorded monkey hand kinematics with eight cameras arranged in a vertical semicircle around the grasp workspace, with denser coverage on the radial side to better capture digit motion and partial views of the torso for later pose analysis. We initially used 1440×1080 FLIR Blackfly S cameras (max 228 fps, BFS-U3-16S2C-CS) and later upgraded to 2048×1536 (max 118 fps, BFS-U3-32S4C-C); lenses were selected to provide sufficient depth of field across the full range of object positions (FLIR LENS-28C2-V100CS, later Computar M0828-MPW3 for better depth of field).

Video streams were recorded across multiple PCs and synchronized by using one camera as a hardware trigger. Frame rates were initially limited by USB3 bandwidth (≈50 fps for the first camera set; ≈40 fps for the higher spatial resolution set). Later, this limit was increased to 90 fps by distributing cameras across four recording PCs. Camera instrinsics were calibrated after installation (and after any lens changes) to correct optical distortion (Greenspon and Sobinov, 2021). Extrinsic calibration was performed before each session using a single-shot Perspective-n-Point procedure (Marchand et al., 2016; OpenCV, n.d.) using a Charuco board placed in view of all cameras to account for day-to-day shifts in camera pose; the board was suspended to provide a stable reference for the vertical axis.

#### Electrophysiological recordings

We recorded single unit and multiunit neural activity using either chronic or acute electrophysiological preparations. Surgical procedures consisted of the implantation of a head-fixing titanium post onto the skull, craniotomy, and implantation of either a recording array (chronic setup) or chamber (acute setup). All surgical procedures were performed under aseptic conditions and anesthesia was induced with ketamine HCl (20 mg per kg, IM) and maintained with isoflurane (10–25 mg per kg per hour, inhaled). Monkeys were handled monkeys in accordance with the rules and regulations of the University of Chicago Animal Care and Use Committee. Monkeys received care from a full-time husbandry staff, and a full-time veterinary staff monitored their health.

Monkeys M and P were each implanted with one 96 channel Utah Electrode array (UEA) with 1.5mm long electrodes, uniformly spaced at 400µm on a 10 by 10 grid. We used anatomical landmarks along with intraoperative surface stimulation of precentral gyrus to target upper limb representations of primary motor cortex. Neural data was collected with CerePlex Direct data acquisition system coupled with CerePlex E digital headstage (Blackrock Microsystems). For each channel, we bandpass filtered neural signals from 250Hz to 3kHz and extracted threshold crossing at a -3.5 RMS value.

Monkey D was implanted with a single recording chamber (Crist Instrument) positioned over primary motor and somatosensory cortices. Stereotaxic coordinates for chamber placement were determined from magnetic resonance imaging (MRI) scans obtained before implantation and anatomical landmarks during implantation. For each session, we placed a microdrive (NAN Instruments) over the chamber, secured one sharpened dura-piercing guide tube in the target location and lowered one Neuropixels probe (Neuropixel 1.0 NHP Long, imec, Belgium) through the guide tubes up to 5 mm below the cortical surface. We used Intan headstages and an OpenEphys acquisition system to amplify and record the signals acquired by the probe. Continuous band-pass filtered (250Hz – 3kHz) signals on each channel were monitored through the OpenEphys GUI (graphical user interface) during probe lowering, stabilization, and recording. We classified the putative units on each electrode as tactile, proprioceptive, or motor based on qualitative assessment of neural responses, using both visual inspection and auditory monitoring during active task performance and passive manipulations, including joint displacements, cutaneous stimulation, and muscle palpation across multiple body regions.

For UEA recordings, we used offline spike sorting software (Offline Sorter, Plexon, Dallas, TX) to remove non-spike threshold crossings and isolate individual units. For Neuropixels probe recordings, we used SpikeInterface (Buccino et al., 2020) for neural signal processing, an open-source Python framework that integrates all steps of the spike-sorting pipeline. This framework was used to remove bad channels, correct for probe drift, detect spikes, and run the spike sorter. We used Kilosort 4 (Pachitariu et al., 2024) through SpikeInterface for automated spike sorting. The resulting outputs were then imported into Phy (Cortex Lab, UCL), a graphical user interface for visualization and manual curation of large-scale electrophysiological data, where they were inspected and curated manually.

#### Experimental control and synchronization of data streams

A single control program coordinated the experiment and communicated with the rest of the setup over TCP/IP. It controlled the robots, target/feedback LEDs, rest-position checks, reward delivery, and auditory cues. Each session was specified by a trial list; failed trials were logged and appended to the list for later repetition. Trials were scored successful when total force fell within the target window and each plate contributed at least 20% of the total, with an option for the experimenter to manually trigger reward to shape behavior. Overall, we collected more than 3500 successful trials of synchronized data in 20 sessions with three monkeys, covering multiple repetitions of the kinematic and force conditions (**Table 1**).

**Table 1.**
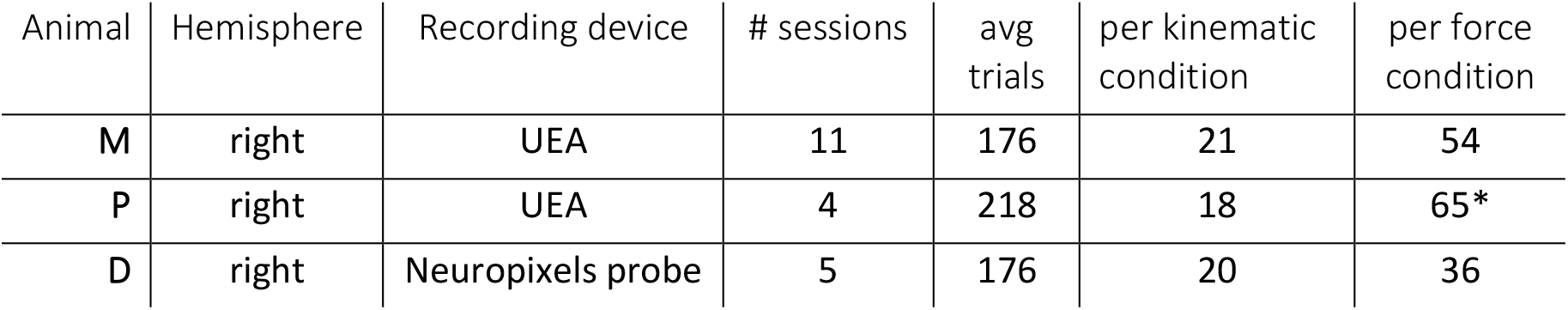
Animals and conditions. Number of trials corresponds to the average number of successful trials per session. Number of trials per condition corresponds to the number of trials on average per session. * some of the sessions with monkey P had only one target force, which were excluded during calculation of number of trials per force conditions.

| Animal | Hemisphere | Recording device | # sessions | avg trials | per kinematic condition | per force condition |
| --- | --- | --- | --- | --- | --- | --- |
| M | right | UEA | 11 | 176 | 21 | 54 |
| P | right | UEA | 4 | 218 | 18 | 65* |
| D | right | Neuropixels probe | 5 | 176 | 20 | 36 |

The control program also interfaced with the camera and pressure-sensor acquisition processes via gRPC messages on the local network (grpc.io, Google, open source). These processes received start/stop recording commands and clock-synchronization messages, which were issued at the start of each trial and paired with a TTL pulse recorded by the neural acquisition system. Round-trip synchronization and acknowledgement were completed within ∼2 ms, and camera processes reported data-save completion at trial end to prevent advancing to the next trial before acquisition finished.

### Data processing pipeline

The recorded data consists of synchronized videos from eight cameras, high-density time-varying pressure sensor maps, and corresponding electrophysiological recordings from the brain. To take full advantage of these data, we computed intrinsic variables defined in body space, as opposed to the external space (**Figure 4**). Those variables – joint angles of the body and forces localized to the segments of the hand – are posited to be more closely related to the representation within primary sensorimotor cortex of behavior and sensory stimuli. We first extracted 3D locations of the skeletal body landmarks using custom trained machine vision networks. Then, we solved the inverse kinematics problem to find joint angles between the segments of the body. Third, we computed which sensels are associated with which hand segment and their relative locations in space, yielding forces applied to each segment. Lastly, we estimated the actuating forces from the corresponding contact locations, measured forces, and the physical properties of the skeletal model.

**Figure 4.**
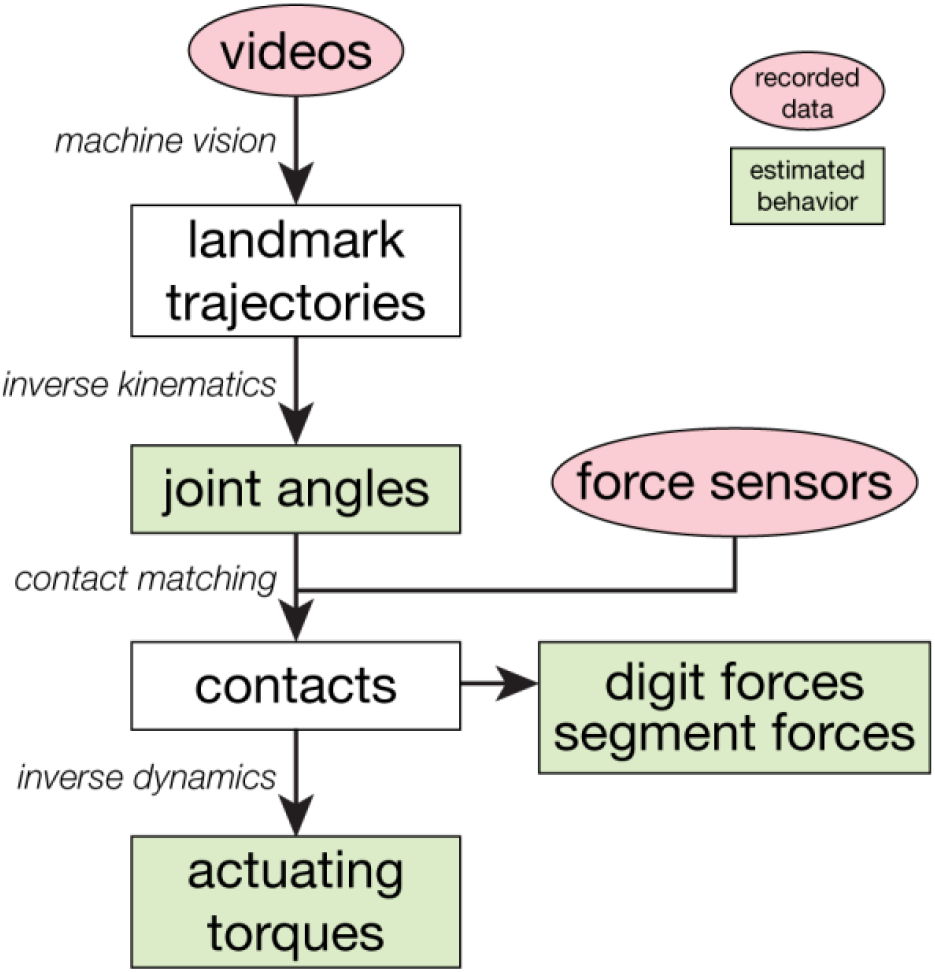
Data processing pipeline. Recorded data marked in red, calculated behavioral data – in green.

#### 3D tracking

We tracked 34 bony landmarks (digits, hand, forearm, arm, torso, **Supplementary Figure 2**), together with the corners of the pressure sensors, initially using DeepLabCut (Mathis et al., 2018), then later a Jarvis-based 2D/3D CNN pipeline (Neurobiology Lab of the German Primate Center, DPZ). A different network was trained for each monkey (and, for Jarvis, per body region: hand, arm, sensors) using 1600-2000 labeled frames. The monkeys were shaved and landmark locations tattooed to improve tracking.

We selected only 2D estimates with likelihood exceeding a threshold (≤0.4 set to NaN) and smoothed/interpolated the resulting trajectories with a 100-ms Gaussian filter. Then, we performed 3D reconstruction using multi-camera triangulation with daily Charuco-based extrinsic calibration. Whenever markers were visible in more than two cameras on a frame, we combined estimates from all pairs of cameras and used centroid of estimates from each pair as the result. Jarvis produced per-frame 3D estimates based on synchronized multi-camera labels, which were smoothed with the same pipeline.

#### Inverse kinematics

3D marker trajectories describe motion in an extrinsic (world) coordinate frame, whereas the motor system controls the limb through muscles, which affect joint angles – an intrinsic frame. The two coordinate systems are related through a kinematic chain, where *forward* kinematics maps joint angles to Cartesian positions, while *inverse* kinematics infers joint angles from measured marker positions. Inverse kinematics is typically an underdetermined problem because of the limb’s redundancy. We solved inverse kinematics with standard optimization (OpenSim), fitting a skeletal model to the 3D markers while enforcing physiological constraints. This reduces noise by using multiple markers per segment and enriches the data with knowledge about anatomy.

Because no publicly available macaque upper-limb model was adequate for our needs, we used a published human upper-body model scaled to each animal’s proportions, as in prior work, accepting the modest anatomical differences between species (Blana et al., 2017; Sobinov, 2022). The model was scaled to each individual using 3D marker locations produced in the previous step and validated against morphometrics measured from the monkeys under anesthesia. Scaling could not readily rely on a standardized calibration pose, as it is done in humans, so we used an iterative procedure that alternated inverse-kinematic fitting and segment-length updates. Starting from a downscaled human model, we selected a time window with maximal marker visibility (typically during grasp), solved inverse kinematics, compared model inter-marker distances to the recorded distances, and scaled each segment accordingly, similar to some automated methods developed for humans (Firouzabadi et al., 2024). We repeated this loop until segment-length changes were <1%, with occasional manual adjustments or exclusion of unreliable markers to avoid poor local minima. Scaling was performed once per animal and was the most labor-intensive step in the pipeline.

We adapted the inverse kinematics pipeline to our task and the scale of movements. To capture both fine digit kinematics and large motion of the hand due to proximal joint rotations, we tightened solver tolerances, which increased occasional frame failures; to avoid aborting trials, we implemented a retry step that temporarily relaxed tolerances for problematic frames. Because the torso was largely stationary, we estimated thorax pose once per session from thorax/clavicle markers and then fixed it to the session mean while solving arm and hand kinematics, reducing spurious thorax and joint motion when proximal markers were missing or noisy. Joint angles, solved frame-by-frame, were then smoothed with a 20 Hz second-order low-pass Butterworth filter followed by a 100 ms Gaussian filter.

To identify the relative location of the pressure sensors to the hand, we modeled each sensel in the same kinematic chain and fit their positions to their marker locations. Sensors containing arrays of sensels were “attached” to the thorax (rather than to the world) so that global reorientation required rotating only a single body - thorax. To reduce the number of outliers, we excluded sensor corner marker points that lay farther from the marker centroid than the sensor’s maximum dimension. During the hold period, sensor pose was treated as rigid and fixed to the median estimate across the interval.

The computed joint angles were consistent with the expected movement patterns. Before the object is grasped, the joint angles reflect the preshaping of the hand to the object shape (**Figure 5**A,B). While the object is being held, hand posture does not change much, beyond a slow drift of a few joints. Varying the kinematic conditions of the task – aperture and tilt of the object – allowed us to increase the effective dimensionality of the kinematics (**Figure 5**C), revealing neurons tuned to both object tilt and aperture (**Figure 5**D). Consistent with previously published findings (Reina et al., 2001), activity of these neurons typically peaked during reach and remained high around grasp onset. Firing rates were generally low during hold period, although many units were dependent on kinematic conditions. We saw units that were tuned to individual kinematic features – e.g., aperture (D221) or tilt (D110) – or several at the same time (D88). Generally, neural activity expressed the complexity expected from previous studies of grasp (Goodman et al., 2019).

**Figure 5.**
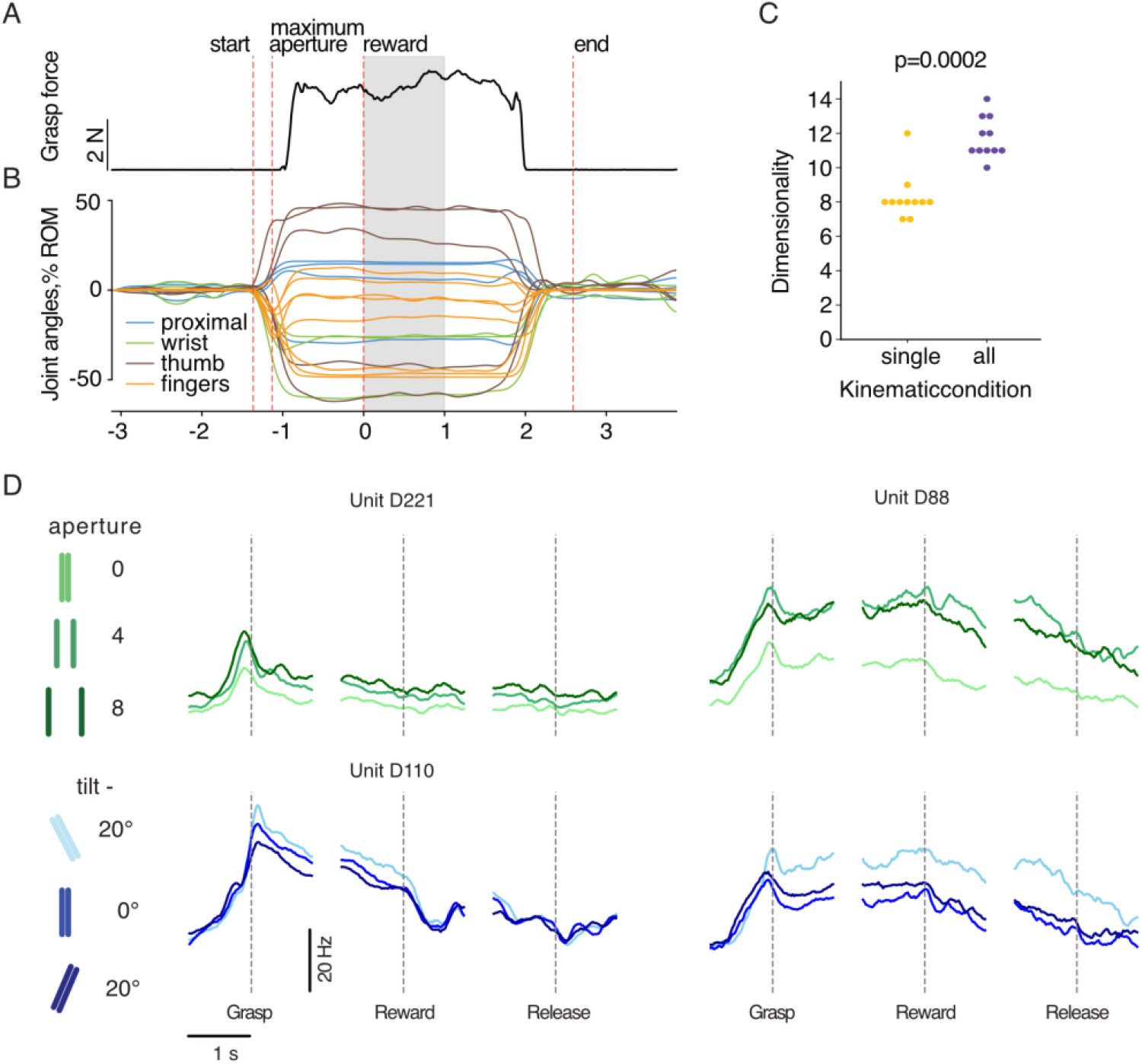
Kinematics. **A**| Total grasp force during a trial. **B**| Example joint angle deviation from starting posture during grasp. Proximal: shoulder and elbow joints; fingers: MCP joint angles. Shaded region denotes the period when the reward was distributed. **C**| Number of principle components to describe 99% of variance, calculated when considering each kinematic condition separately, or all together. Mann-Whitney U=114. **D**| PETHs of several individual motor cortical neurons from monkey D averaged over kinematic conditions and aligned to grasp and reward.

#### Mapping recorded forces to hand locations

To assign force measurements to hand segments, we converted the scaled arm, hand, and object models in OpenSim to MuJoCo while preserving joint locations and geometry (Todorov et al., 2012). The object surfaces were represented as sensel grids. For efficiency, we retained only sensels with nonzero values in a session (a reduction from 1936 to ∼400–800 per plate) and downsampled to the camera frame rate using a median filter to create synchronized kinematic-force frames.

We then replayed grasp kinematics in MuJoCo and, at each time point, assigned each active sensel to the nearest hand segment, such as the distal phalanx of the middle finger, or proximal phalanx of the thumb, generating a time series of segment-sensel contact pairs (**Figure 6**A). Segment forces were computed by summing forces across assigned sensels. After all the contacts were calculated, we up-sampled the digit force back to the rate of the force recording – 300 Hz. Contact assignments were interpolated between kinematic frames, switching attribution at midpoints when contacts appeared, disappeared, or shifted between segments. The spatial resolution of these signals (1.9x1.9 mm^2^) is lower than most cutaneous receptive fields in glabrous skin (Johansson and Flanagan, 2008) but still present a high-resolution look at the forces encountered by the skin during natural tactile interaction.

**Figure 6.**
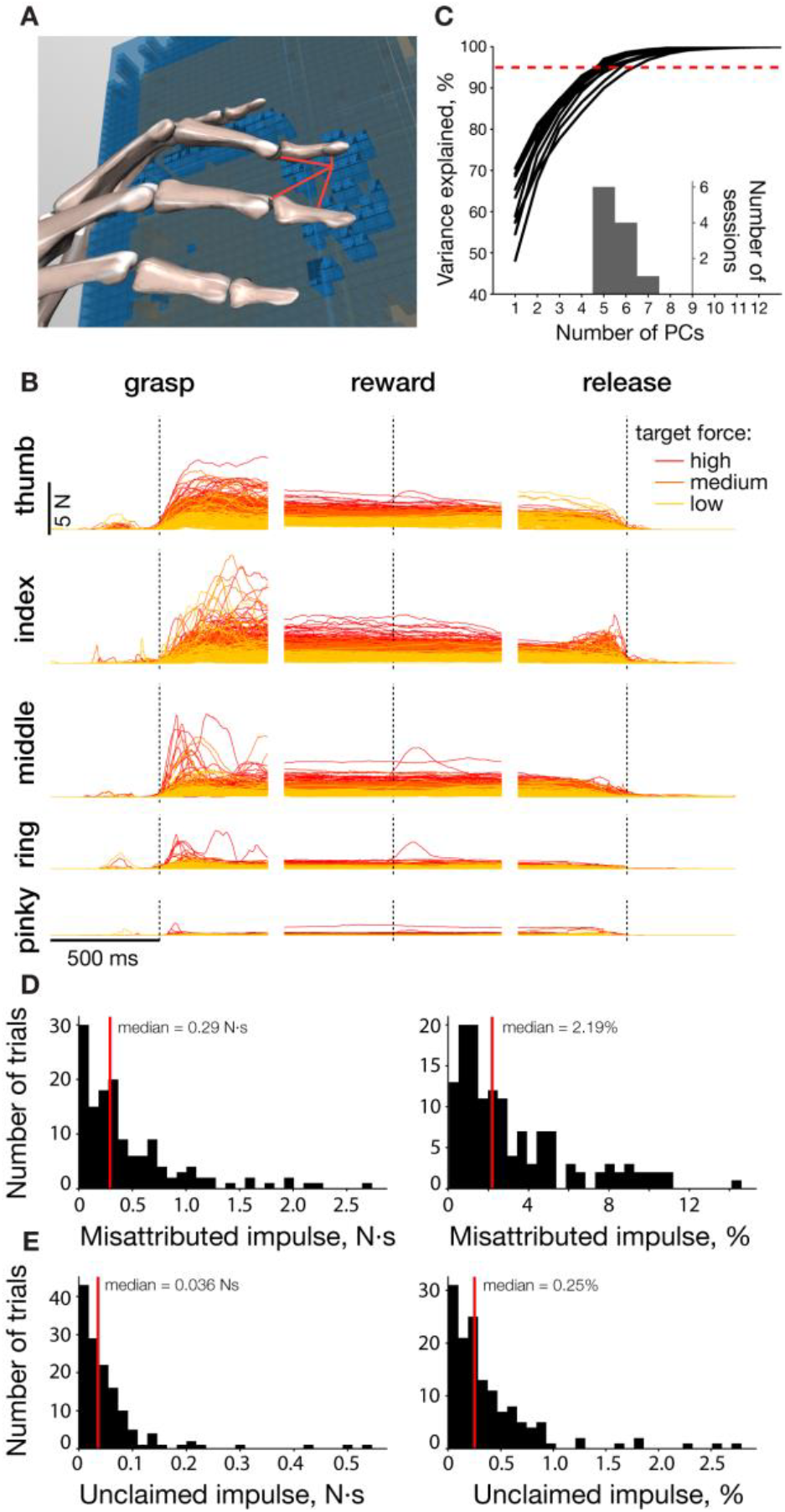
Mapping the pressure to the hand. **A**| Models of the hand and pressure sensors in MuJoCo. Sensels are shown as blue boxes. Sensels with non-zero force are opaque, and otherwise, transparent. Red lines show example hand segments to which the distances were calculated from a single sensel. **B**| Forces applied by each finger in every trial of a single session, color indicates the target force. **C**| Cumulative variance of force separated by segments explained by the top principal components (PCs). Dashed red line indicates 95%. Insert: Number of PCs necessary to explain 95% of variance. **D**| Distribution of impulse attributed to the wrong segment or no segment, absolute (left) and normalized (right) for monkey M over 138 trials in 11 experimental days. Red line denotes median. **E**| Same as D for impulse unclaimed during automatic matching process.

The force profiles for each digit were broadly proportional to the total grasp force – the traces for high grasp force higher than those for low grasp forces (**Figure 6**B). However, the forces applied by each digit segment were not perfectly correlated with each other. To quantify the degree of independence, we performed principal component analysis on segment forces for all sessions of monkey M (**Figure 6**C). The number of PCs needed to describe 95% of the variance across all 19 hand segments varied between 5 and 7, highlighting the multi-dimensional nature of the grasp force.

We validated automatic force attribution against manual pressure maps generated by experimenters for monkey M, using the following process. Using sensel pressure maps synchronized to video, an evaluator assigned sensels to digits for a number of randomly selected trials from each session. We quantified a “disagreement” metric as the time integral of the difference between manually and automatically attributed forces. The metric calculated misattributed impulse (N·s) per trial, which we normalized by the total impulse. Median error was around 2% in monkey M (corresponding to about 0.29 N·s) over the whole trial (**Figure 6**D), indicating close agreement between automatic and manual labels. We also quantified force that was not assigned to any segment as “unclaimed” impulse and found it to be an order of magnitude smaller (median 0.036 N·s, 0.25%). Together, these metrics indicate the high precision of the automated force-allocation procedure.

#### Inverse dynamics

The final building block in the construction of the full physical description of grasping is the estimation of actuating torques generated by the muscles that produce both the movement of the hand and the forces applied to the object. Estimation of torque requires knowing the kinematics of the movement – posture, velocity, and acceleration of joints – and definition of the forces acting in the system. There are three major forces: gravitational, interaction (describing attachments between segments and their effect on each other), and other external (forces of interaction with the object). The mass of each segment was obtained by scaling that of the human model to the size of each monkey. Interaction forces were computed by the MuJoCo physics engine (Todorov et al., 2012) using the defined kinematic chains. We estimated the joint damping coefficients based on the published literature in humans for digits(Fiorilla et al., 2011; Binder-Markey and Murray, 2017; McFarland et al., 2023; Tsakonas et al., 2024), wrist(Park et al., 2011; Klomp et al., 2018; Falzarano et al., 2021), elbow(Popescu et al., 2003; Selen et al., 2007), and shoulder(Lipps et al., 2020; Yahya et al., 2022). The last part – interaction forces – we computed as described above. Here, we applied them as external forces acting along the normal axis to the surface of the sensel, matching how the forces were measured (**Figure 7**A). Given all three components, torques were calculated at each time point (**Figure 7**B).

**Figure 7.**
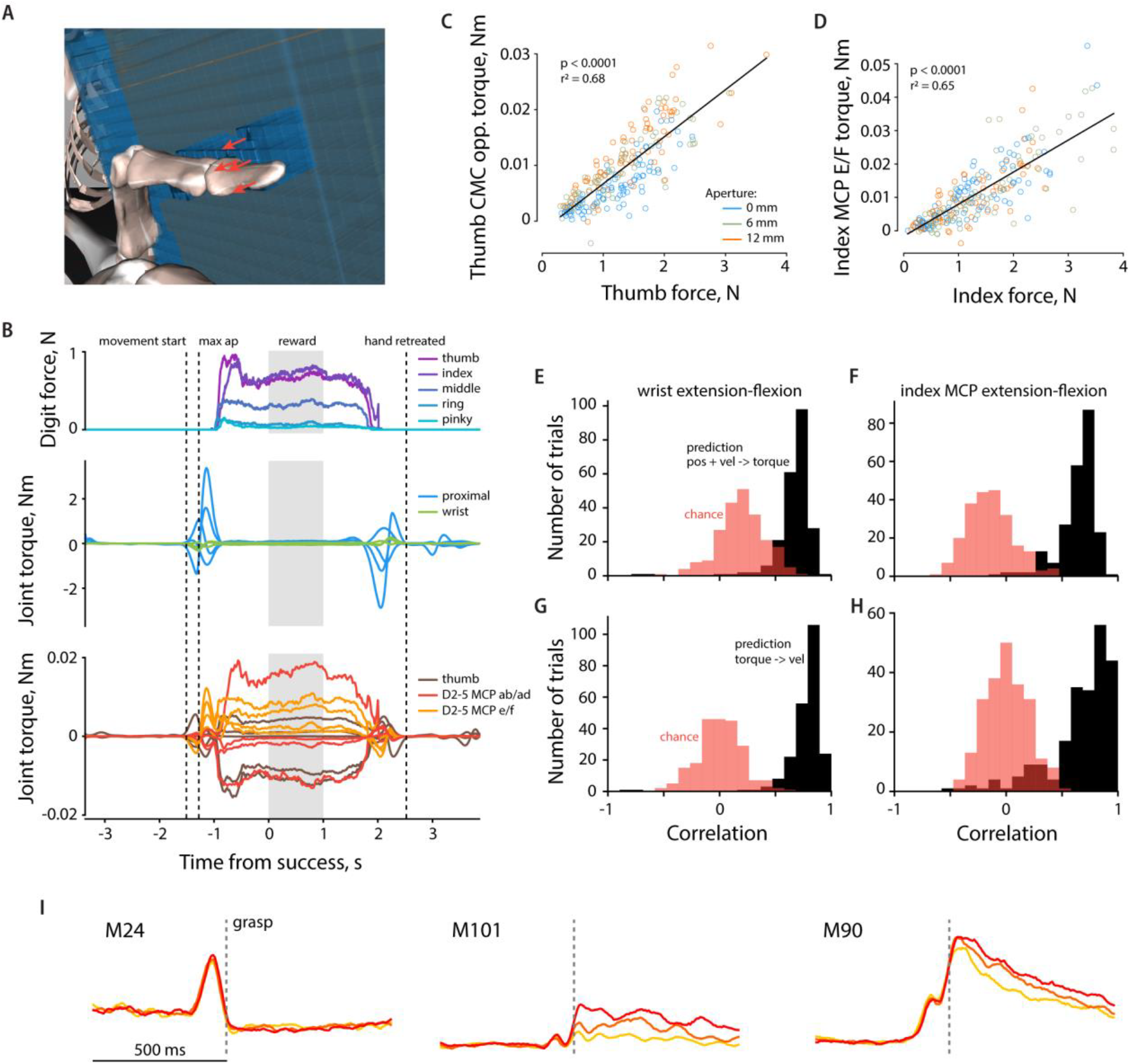
Actuating torques. **A**| Schematic of the modelled external forces applied to a single segment – the distal phalanx of index finger. Red arrows represent the direction of the external forces exerted by the sensels on the distal index segment. The length of the arrow does not represent force magnitude. **B**| Forces and torques of select joints during grasp. **C**| Relationship between thumb CMC opposition torque and the amount of force exerted by the thumb. **D**| Same as C for index MCP extension-flexion and force. **E**| Pearson correlation between wrist flexion-extension torque computed from dynamics and torque predicted by cross-validated ridge regression from wrist position and velocity. Red: time-shuffled prediction. **F**| Same as E for index MCP extension-flexion joint. **G**| Pearson correlation between wrist e/f velocity and velocity predicted by cross-validated ridge regression from wrist e/f torque. **H**| Same as G for index MCP e/f. **I**| Example single unit PSTHs aligned to grasp onset, averaged within grasp force target and color coded by the grasp force. Left: proximal-like, middle: distal-like, right: mixed. MCP: metacarpophalangeal; CMC: carpometacarpal; ab/ad: abduction-adduction; e/f: extension-flexion; opp.: opposition.

Measuring actuating torques during movement for validation is nearly impossible, as it requires insertion of force-measuring devices into and around joints(Ravary et al., 2004; Bergmann et al., 2007, 2014; Go et al., 2017; Zhang et al., 2021; Escriche-Escuder et al., 2022; Rastegarpanah and Taylor, 2024). Our calculated torques, however, match the ranges previously reported in the modeling literature for individual MCP joints during free movement and light object interaction(Jindrich et al., 2004; Grinyagin et al., 2005; Balakrishnan et al., 2006; Nataraj and Li, 2015) (**Figure 7**BC). Furthermore, the time courses of estimated torques fit the expected qualitative observations. First, torques at the proximal joints were large during the reach phase, as the monkey lifted its arm and forearm to reach toward the object, and during return to the resting posture (**Figure 7**B). When the monkey started grasping the object, digit torques rose to produce the grasping forces (**Figure 7**B, top). During the grasp, the torques were higher than during the preshaping, remained relatively constant, and were closely related to the force exerted by the respective digit (**Figure 7**CD). However, the relationship between force and torque was affected by kinematic conditions (**Figure 7**C). Going from small object width (blue circles, below the regression line) through medium (green) to large (orange, above the regression) width, the thumb opposition torque increased, matching the change in the grasping strategy monkey was employing. This relationship was dependent on the joint and, for example, not present in the index finger extension/flexion(**Figure 7**D).

During grasp, torques primarily reflected force-related features, while during preshaping they reflected kinematics. For two example joints, wrist extension-flexion and index MCP e/f, we linearly predicted joint torque during preshaping from the joints’ angular position and acceleration, with accuracy well above models built on time-shuffled data (**Figure 7**EF). The relationship was bidirectional, as torque could also be used to predict velocity in the same joint (**Figure 7**GH), further underlining the close relationship between kinematics and torques during free hand movements. However, these predictions did not capture all variability, indicating that joint kinematics and torques provide related but nonredundant descriptions of behavior. Kinematics describe how the hand moves, whereas torques capture the mechanical demands required to generate and control that movement. Thus, a complete physical account of manipulation requires considering a combination of both signals.

As was the case for the kinematics-related neuornal examples shown in Figure 5, we observed diverse single-unit responses that qualitatively tracked torque dynamics (**Figure 7**I). Some resembled proximal joint torques, with large transients during reaching and no separation across force levels during the hold (**Figure 7**I, M24). Others resembled torques at distal joints, exhibiting clear force-dependent modulation after contact with minimal pre-contact activity (**Figure 7**I M101). A third group showed mixed patterns consistent with contributions from both proximal and distal torques (**Figure 7I** M90), possibly related to muscles that span multiple joints (e.g., the long finger flexors). Together, these response types suggest that M1 activity can reflect the mechanical demands of interaction across multiple joints.

## Discussion

We have described in detail an approach for quantifying both the kinematics and kinetics of object grasp. This combination of detailed motion estimates and high-resolution contact-force measurements opens the door to analyzing how neural activity represents object interaction and manipulation in otherwise inaccessible, intrinsic coordinates.

The contact force data provide access to the continuous cutaneous signals affecting the somatosensory system during grasp. Because these measurements operate at a fine spatial scale and capture rapid contact transients, they are well suited to linking contact mechanics to neural responses (Vega-Bermudez and Johnson, 1999; Saal et al., 2017). Previous work has characterized receptive-field structure and temporal response properties primarily using passive or otherwise tightly controlled tactile stimulation (DiCarlo et al., 1998; Delhaye et al., 2018; Long et al., 2022). Our approach extends that line of work into active exploration and manipulation, where sensory feedback is continuously integrated with ongoing movement (Chapman and Ageranioti-Bélanger, 1991; Johansson and Flanagan, 2008, 2009; Simões-Franklin et al., 2010; Omrani et al., 2016).

Kinematics are informative during the transport and hand preshaping phases but often become less useful after contact is established. In contrast, contact forces are only present during interaction with the object. Estimating joint torques provides a way to describe both preshaping and grasping within a shared mechanical framework. Combined with kinematics, torques capture important aspects of the limb’s physical properties and its interaction with the object. Such variables may be particularly relevant if neural activity reflects internal models of movement dynamics(Wolpert et al., 1995; Hwang and Shadmehr, 2005; McNamee and Wolpert, 2019). Torques are also closely tied to motor output, as they emerge from muscle forces filtered through the mechanics of the musculoskeletal system, though they do not uniquely determine individual muscle activations (Flash and Mussa-Ivaldi, 1990; Simpson et al., 2015; Olesh et al., 2017; Mathieu et al., 2023).

On the sensory side, torque estimates open an opportunity to consider force-related proprioceptive signals associated with Golgi tendon organs. Although Golgi tendon organs are less frequently emphasized than muscle spindles, they constitute a substantial portion of the proprioceptive afferents and carry information about muscle load (Prochazka, 2015; Sobinov and Bensmaia, 2021). By contrast, muscle spindle signals are more closely related to muscle length and its rate of change and, can be approximated with musculoskeletal models that reconstruct muscle paths and stretch (Sartori et al., 2012; Sobinov et al., 2020; McFarland et al., 2023). However, predicting spindle afferent activity during active behavior remains difficult without additional information about muscle activation, because spindle responses depend not only on muscle mechanics but also on descending fusimotor control (Prochazka, 1999; Blum et al., 2020; Dimitriou, 2022).

The pipeline described here makes use of several technologies across different processing steps. While we made particular choices for implementation, the overall logic is modular and could be reproduced with alternative tools that provide equivalent outputs. We used two markerless vision-based approaches for tracking body landmarks, but this area is evolving rapidly. Many public models are now available for human pose estimation, although not for other primates (MMPose Contributors, 2020; Falisse et al., 2025; Mundt et al., 2025). A key limitation, however, is that these models often predict generalized keypoints near joint centers rather than anatomically defined skeletal landmarks. Such points define approximate ball joint locations, but they do not necessarily provide stable markers of rigid body segment pose. Because they may shift relative to the underlying segment across postures or imaging conditions, their use in inverse kinematics can introduce instability and should be treated with caution. Progress in markerless tracking for nonhuman primates has also been substantial, but performance and generalization still lags behind human pose estimation because datasets are magnitudes smaller and less standardized, especially for the hands (Hayden et al., 2022; Yao et al., 2023). For this reason, marker-based infrared motion capture remains an important option when very high spatial precision is needed, despite the added burden of setup and repeated marker placement (Scataglini et al., 2025).

We also treated 3D landmark estimation and inverse kinematics as separate steps, although recent work increasingly integrates these operations. Some methods augment sparse video keypoints into dense anatomical markers before running inverse kinematics, whereas others directly estimate joint angles with a skeletal model embedded into the learning process (Firouzabadi et al., 2024). Such approaches can improve anatomical consistency and reduce the impact of occlusions or noisy detections.

We placed pressure sensors on the objects rather than on the hand. Although glove-based sensors would simplify contact localization and support a wider range of object shapes, current hand-shaped sensors have lower sensel density, can interfere with natural movement, and in our experience reduce the accuracy of hand-tracking algorithms. They are also far more difficult to use with monkeys than humans. Object-mounted sensors preserve more natural behavior but constrain the shapes that can be tested, because sensors cannot be reshaped without recalibration. This limitation could be addressed by preparing and calibrating multiple sensorized objects offline and swapping them during experiments.

There are several natural extensions of this apparatus that could address additional questions. For example, the object surface could be covered with materials with different textures, including frictional properties. Although very compliant materials would likely interfere with force measurements, rigid low and high friction surfaces could be incorporated readily. This would be useful because friction is a major determinant of grip-force scaling during object manipulation, and friction- and slip-related tactile signals strongly influence both manipulation behavior and somatosensory responses (Cadoret and Smith, 1996; O’Shea and Redmond, 2021). In these experiments, the subject would have to pull or push the object to produce tangential friction forces. Because the object is robotically controlled, it could also be perturbed after contact to simulate transport, in-hand rotation, or unexpected changes in pose. If the objects were hidden before reach onset, the perturbations of object shape and location could be used to study prediction and expectation during grasp (Castiello et al., 1998). Likewise, unexpected changes in shape or load could probe rapid corrective responses, extending perturbation paradigms that have so far been examined mostly in reaching and more constrained grasping tasks (Picard and Smith, 1992; Roy et al., 2006; Pruszynski and Scott, 2012; Camponogara and Volcic, 2019).

This framework is adaptable to smaller model organisms, including rodents, extending established rodent reach-to-grasp paradigms to more direct measurements of contact forces during grasp. Doing so would likely require custom miniaturized sensor arrays and object shapes tailored to the animal and task, but the overall logic of the pipeline would remain similar. Such an extension appears feasible because rodent skilled reaching behavior is well established (Bova et al., 2019; Forghani et al., 2023; Grier et al., 2026), and musculoskeletal models of the mouse forelimb and whole body are available (Tata Ramalingasetty et al., 2021; Gilmer et al., 2025). However, direct measurement and biomechanical reconstruction of grasp remain limited. Applying this approach in rodents would provide a deeper understanding of grasping in a model organism that also has available, powerful genetic tools.

## Author contributions

ARS and SJB designed the study with input from other authors.

XM, EVO, QH, NS performed the experiments.

ARS, CMG, CR, PJ, PA constructed and programmed the experimental devices.

ARS designed the data processing pipeline.

ARS, XM, CMG, CR, ND, RB developed aspects of data processing.

ARS wrote the first draft of the manuscript.

ARS and XM wrote the paper with input from all authors.

## Competing interests

ARS was a consultant for MuJoCo (DeepMind, Google) on the VR implementation during this study. CMG has received sponsored travel from Blackrock Neurotech. NGH serves as a consultant for Blackrock Neurotech.

Acknowledgements

The authors would like to thank the veterinary staff at the University of Chicago Animal Research Center for their help. This work was supported by the National Institute of Neurological Disorders and Stroke, R35 NS122333, R01 NS125270.

## Data availability

A sample session with all data with monkey M will be made available online upon publication.

## Code availability

All code required to run the experiments, process the data, and perform the analysis are going to be made available in public repositories:

https://github.com/SobinovLab/prehension_methods_paper – manuscript figures and example scripts. https://github.com/SobinovLab/prehension – data processing and analysis.

https://github.com/SobinovLab/stereognosis_protocol_main – main protocol for conducting experiments and controlling robots.

https://github.com/SobinovLab/tekscan_pressure_sensor_server – recording data from pressure sensors while providing feedback.

https://github.com/SobinovLab/cameras_server – recording synchronized videos from high-speed FLIR cameras.

## Supplementary materials

**Supplementary Figure 1.**
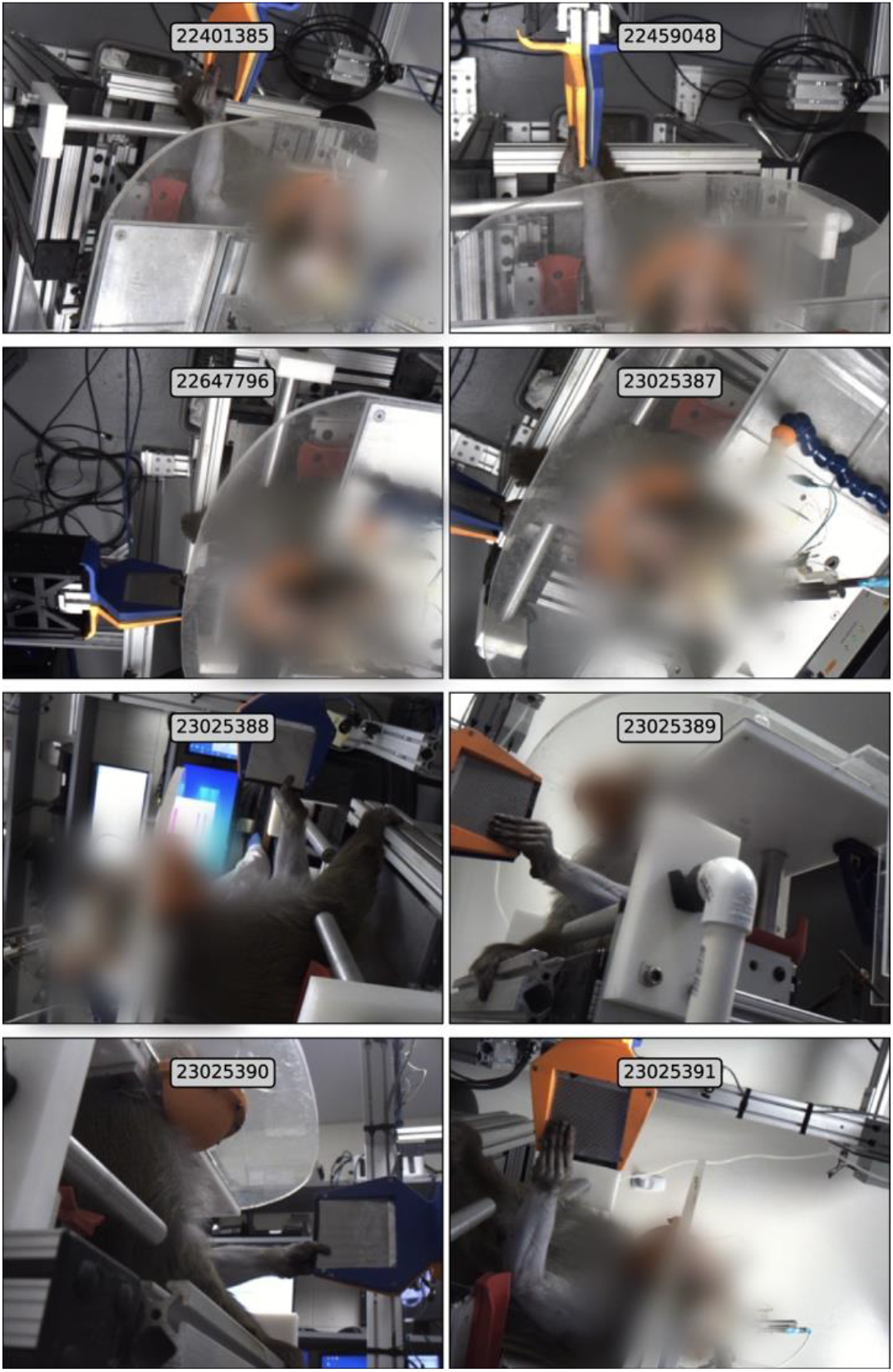
Views from the eight cameras during object grasp at the moment of successful grasp. The numbers correspond to the camera serial numbers.

**Supplementary Figure 2.**
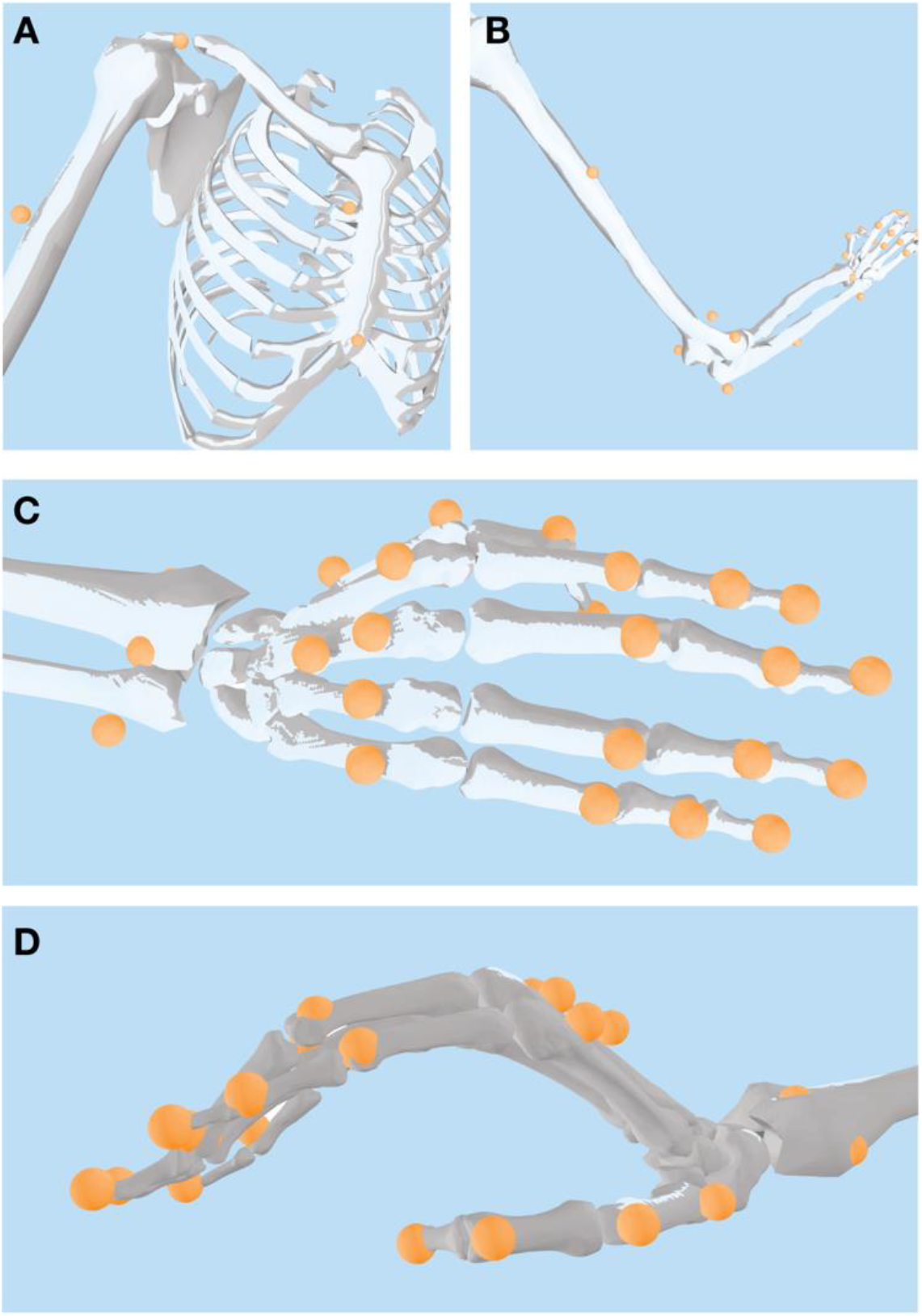
Locations of markers on a primate skeleton used as a reference during training of the network. The shown OpenSim model was taken from (Sobinov AAH, 2022) and scaled to monkey M. Upper sternum marker has been shifted right to match the location of the tattoo.

**Supplementary Figure 3.**
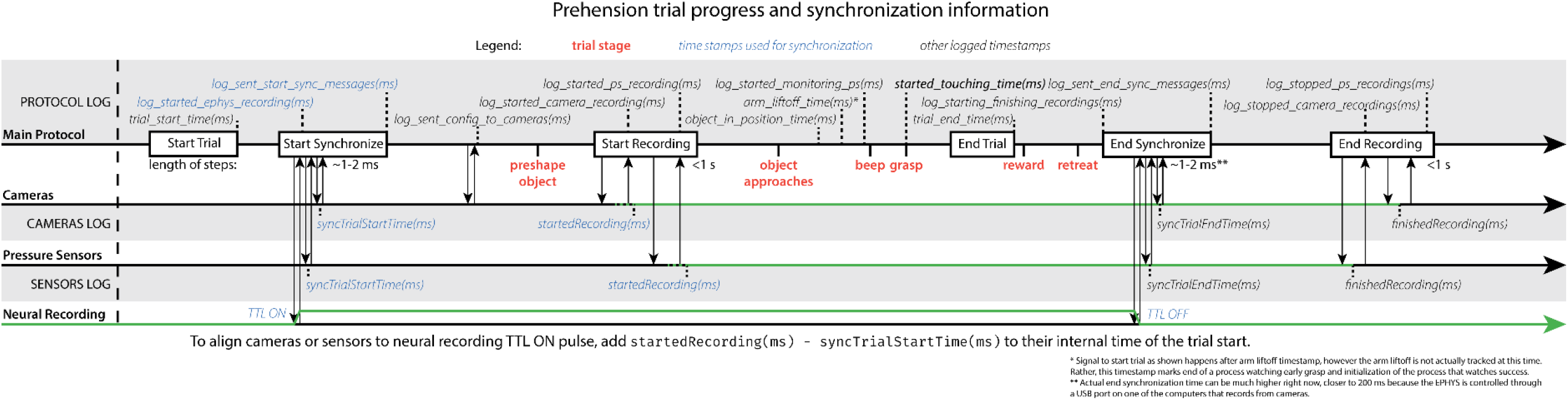
Communication and messages between recording and control modules.

**Supplementary Figure 4.**
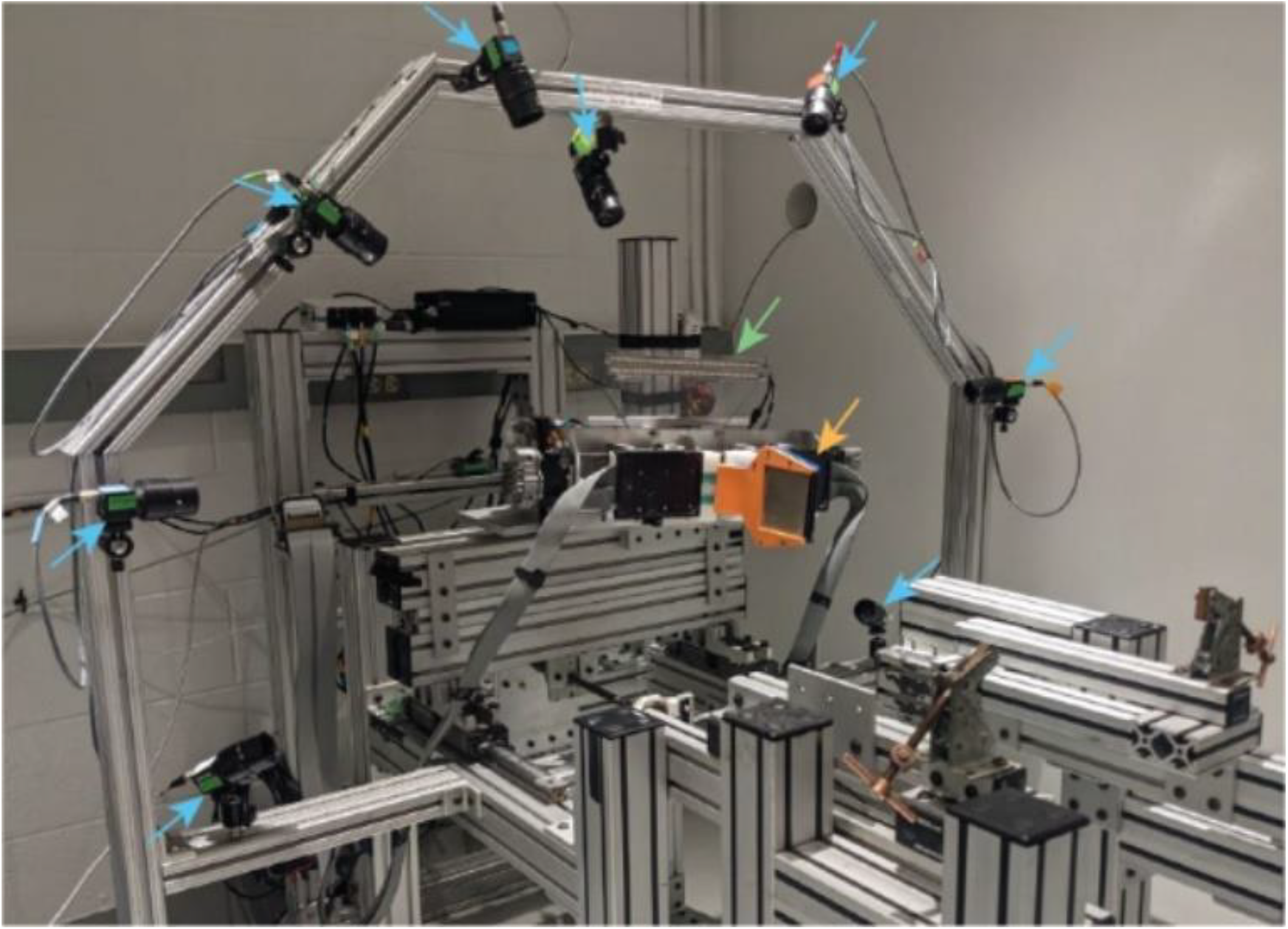
Experimental rig actuated by servomotors.

